# Reward boosts reinforcement-based motor learning

**DOI:** 10.1101/2021.03.23.436627

**Authors:** Pierre Vassiliadis, Gerard Derosiere, Cecile Dubuc, Aegryan Lete, Frederic Crevecoeur, Friedhelm C Hummel, Julie Duque

**Affiliations:** Institute of Neuroscience, Université Catholique de Louvain, 1200, Brussels, Belgium; Defitech Chair for Clinical Neuroengineering, Center for Neuroprosthetics (CNP) and Brain Mind Institute (BMI), Swiss Federal Institute of Technology (EPFL), 1202, Geneva, Switzerland; Institute of Information and Communication Technologies, Electronics and Applied Mathematics, Université Catholique de Louvain, Louvain-la-Neuve 1348, Belgium; Defitech Chair for Clinical Neuroengineering, Center for Neuroprosthetics (CNP) and Brain Mind Institute (BMI), Swiss Federal Institute of Technology Sion (EPFL), 1951, Sion, Switzerland; Clinical Neuroscience, University of Geneva Medical School (HUG), 1202, Geneva, Switzerland

**Keywords:** Motor learning, motor skill learning, reward, motivation, reinforcement, learning, sensory feedback, motor control

## Abstract

Besides relying heavily on sensory and reinforcement feedback, motor skill learning may also depend on the level of motivation experienced during training. Yet, how motivation by reward modulates motor learning remains unclear. In 90 healthy subjects, we investigated the net effect of motivation by reward on motor learning while controlling for the sensory and reinforcement feedback received by the participants. Reward improved motor skill learning beyond performance-based reinforcement feedback. Importantly, the beneficial effect of reward involved a specific potentiation of reinforcement-related adjustments in motor commands, which concerned primarily the most relevant motor component for task success and persisted on the following day in the absence of reward. We propose that the long-lasting effects of motivation on motor learning may entail a form of associative learning resulting from the repetitive pairing of the reinforcement feedback and reward during training, a mechanism that may be exploited in future rehabilitation protocols.

## 1. Introduction

Motor skill learning is the process by which the speed and accuracy of movements improve with practice (Krakauer et al., 2019). It is crucial for daily-life activities (*e.g.*, when learning to tie shoelaces) and motor recovery after injury (*e.g.*, after a stroke, Roemmich and Bastian, 2018). A large amount of research has since long demonstrated that motor learning relies on sensory feedback, which allows the reduction of movement errors (*e.g.*, Shadmehr et al., 2010; Tseng et al., 2007). More recently, some studies have shown that reinforcement feedback, allowing the adjustment of movements based on knowledge of performance, also plays a role in motor learning (Bernardi et al., 2015; Galea et al., 2015; Mawase et al., 2017; Palminteri et al., 2011; Therrien et al., 2016; Wachter et al., 2009). The contribution of reinforcement feedback to motor learning seems to be particularly important when the quality of the available sensory feedback is low (Cashaback et al., 2017; Izawa and Shadmehr, 2011). These observations suggest that reinforcement feedback may be critical for motor rehabilitation (Quattrocchi et al., 2017; Roemmich and Bastian, 2018), where patients often exhibit sensory impairments in addition to their motor disability (Connell, 2008; Hepworth et al., 2016). However, before clinical translation can occur, significant research is required to characterize the optimal conditions in which sensory and reinforcement feedback can improve motor learning.

One key factor that may influence sensory- and reinforcement-based motor learning is motivation (Lewthwaite and Wulf, 2017). This idea is in line with an ethological perspective: in nature, animals are motivated to learn efficiently motor behaviors that have been repetitively associated with rewarding outcomes, in order to increase the likelihood to reach these outcomes again in the future (Barron et al., 2010; Yamazaki et al., 2016). Whereas past research on motivation has traditionally focused on the impact of reward on decision-making (Balleine and O’Doherty, 2010; Bush et al., 2002; Dayan and Niv, 2008; Derosiere et al., 2017b, 2017a; Gershman and Daw, 2017; Hare et al., 2011; O’Doherty, 2004; Padoa-Schioppa, 2011; Schultz, 2015; Shima and Tanji, 1998), there has been a recent rise in interest regarding its influence on motor control (Carroll et al., 2019; Grogan et al., 2020; Manohar et al., 2019; Reppert et al., 2015; Shadmehr et al., 2019; Uehara et al., 2019; Vassiliadis et al., 2019; Yoon et al., 2019). As such, recent studies have shown that the presence of reward can improve both movement speed and accuracy, sometimes even concomitantly (Codol et al., 2020; Manohar et al., 2015; Vassiliadis and Derosiere, 2020). An important question is whether such motivation-related gains in motor control can persist after training, when the reward is removed (Chen et al., 2017; Sporn et al., 2020).

To tackle this issue, previous studies have investigated motor skill learning with different types of reinforcement and reward. This research showed that the combination of reinforcement (providing knowledge of performance) and reward (providing motivation) can influence motor skill learning (*e.g.*, Abe et al., 2011; Steel et al., 2019, 2016; Wachter et al., 2009; Wilkinson et al., 2015). A key aspect of the aforementioned studies is that they considered reinforcement and reward in a bonded way, with the rewarded participants being also the ones receiving performance-based reinforcement feedback. The assumption underlying this approach is that receiving knowledge of performance (*e.g.*, points or binary feedback) provides a form of intrinsic reward that by itself increases motivation to perform well (Leow et al., 2018). However, in addition to the intrinsically rewarding properties of reinforcement, knowledge of performance also provides a learning signal to the motor system, that can influence motor learning (Bernardi et al., 2015; Galea et al., 2015; Huang et al., 2011; Kim et al., 2019; Leow et al., 2018; Mawase et al., 2017; Nikooyan et al., 2015; Shmuelof et al., 2012; Therrien et al., 2016; Uehara et al., 2018). In contrast, extrinsic reward increases motivation to perform well, without conveying any additional learning signal (Berke, 2018). Consistent with a dissociable role of reinforcement and reward in motor learning, previous research has shown that certain subpopulations of neurons in the motor cortex (*i.e.*, a key region of the motor learning network; Krakauer et al., 2019) are responsive to the outcome of previous movements irrespective of reward (Levy et al., 2020), while others respond to reward regardless of the previous outcome (Ramkumar et al., 2016). Put together, these elements indicate that estimating the net impact of reward on motor learning requires controlling for the effect of the reinforcement feedback on the learning process. In the present study, we aimed at studying the specific contribution of reward to motor skill learning and maintenance by experimentally uncoupling it from knowledge of performance.

Another important question relates to how, at the single-trial level, motivation by reward may affect motor skill learning and maintenance. As such, computational models of motor learning posit that movement errors can be corrected based on sensory and reinforcement feedbacks on a trial-by-trial basis (Cashaback et al., 2017), with possible interactions between these two processes (Izawa and Shadmehr, 2011). Sensory-based motor learning relies on the ability to produce motor commands that match target sensory consequences (*e.g.*, visual, somatosensory consequences; Sidarta et al., 2016). Most previous research on sensory-based motor learning has focused on adaptation paradigms where participants have to counter an external perturbation through sensory prediction errors (*i.e.*, corresponding to the difference between expected and received sensory feedback; (Avraham et al., 2020a; Morehead and Orban De Xivry, 2021; Shadmehr et al., 2010; Tsay et al., 2021). In contrast, motor skill learning often requires to improve motor commands to match a target sensory state without any external perturbation (*e.g.*, when learning to play tennis; Krakauer et al., 2019; Sidarta et al., 2016). Here, learning relies on the progressive adjustment of motor commands towards the target sensory state based on previous sensory experience (Bernardi et al., 2015). Conversely, reinforcement-based motor learning is thought to depend on the ability to efficiently regulate between-trial motor variability based on previous outcomes (Dhawale and Smith, 2017; Pekny et al., 2015; Sidarta et al., 2016; Therrien et al., 2016; Uehara et al., 2019; Wu et al., 2014). Importantly, in this framework, reward may have a global effect, enhancing both sensory- and reinforcement-based adjustments from one trial to another, or could have a more specific effect, boosting only one of the two learning systems (Galea et al., 2015; Kim et al., 2019). Here, we sought to investigate the effect of reward on sensory-based and reinforcement-based adjustments in motor commands during motor skill learning at the single-trial level, in a situation where they would both contribute to the learning process.

Healthy participants (n = 90) trained on a pinch-grip force task with limited sensory feedback over two consecutive days, while we manipulated the reinforcement feedback and reward on Day 1. By removing visual feedback on most trials, we ensured that the learning process would largely depend on the integration of somatosensory and reinforcement feedbacks (Bernardi et al., 2015; Izawa and Shadmehr, 2011; Sidarta et al., 2019). Moreover, subjects were distributed in three groups where training involved sensory (S) feedback only (Group_-S_; n = 30), sensory and reinforcement (SR) feedback (Group_-SR_; n=30), or both feedbacks and a monetary reward (SRR, Group_-SRR_; n=30). We investigated how participants learned and maintained the skill depending on the type of feedback experienced during training. We found that while sensory and reinforcement feedbacks were not sufficient for the participants to learn the task in the present study, adding reward during training markedly improved motor performance. Reward-related gains in motor learning were maintained on Day 2, even if subjects where no longer receiving a reward on that day. Importantly, single-trial analyses showed that reward specifically increased reinforcement-related adjustments in motor commands, with this effect being maintained on Day 2, in the absence of reward. The pinch-grip force task used here also allowed to consider adjustments separately for the speed of force initiation, and the accuracy of the performed force, both in terms of variability and amplitude. Importantly, we found that reward did not affect the control of all motor components in the same way, with the amplitude component turning out to be the more strongly influenced by the presence of reward.

Altogether, the present results provide evidence that motivation by reward can improve motor skill learning and maintenance even when the task is performed with the same knowledge of performance. More importantly, this effect seems to entail a specific potentiation of reinforcement-related adjustments in the motor command at the single-trial level. We propose that the long-lasting effects of motivation on motor learning may involve a form of associative learning resulting from the repetitive pairing of the reinforcement feedback and reward during training. These behavioral results are important to characterize the mechanisms by which reward can enhance motor learning and may guide future motivational interventions for rehabilitation (McGrane et al., 2015).

## 2. Results

Ninety healthy participants practiced a pinch-grip force task over two consecutive days. The task required participants to squeeze a pinch grip transducer to move a cursor as quickly as possible from an initial position to a fixed target and maintain it there for the rest of the task (**Figure 1A**). The force required to reach the target (Target_Force_) corresponded to 10 % of the individual maximum voluntary contraction (MVC). In most of the trials (90 %), participants practiced the task with very limited sensory feedback: the cursor disappeared when the generated force reached half of the Target_Force_. In the remaining trials (10 %; not considered in the analyses), full vision of the cursor allowed participants to be visually guided towards the Target_Force_ and therefore to be reminded of the somatosensory sensation corresponding to the Target_Force_. Hence, in this task, learning relied mostly on the successful reproduction of the Target_Force_ based on somatosensory feedback (Raspopovic et al., 2014), with the target somatosensory sensation being regularly reminded to the participants through the full vision trials. To learn the task, subjects were provided with six training blocks (40 trials each; *i.e.*, total of 240 training trials; **Figure 1B**), during which Group_-S_ subjects trained with sensory feedback only (Block_-S_), Group_-SR_ subjects trained with sensory and reinforcement feedback (Block_-SR_), and Group_-SRR_ subjects trained with both feedbacks and a monetary reward (Block_-SRR_). Notably, the groups were comparable for a variety of features including age, gender, Target_Force_, difficulty of the task, muscular fatigue and final monetary gains (see Materials and Methods, **Table 1**). Beside the training blocks, all participants performed the task in a Block_-SR_ setting so that the familiarization, the pre- and post-training assessments on Day 1, as well as Re-test on Day 2, occurred in the same conditions in the three experimental groups. This design allowed us to investigate the effect of reinforcement and reward, both on learning and on maintenance of the learned motor skill. Importantly, in Block_-SR_ and Block_-SRR_, the binary reinforcement feedback depended on the Error, estimated as the absolute difference between the Target_Force_ and the exerted force over the whole trial (excluding the first 150 ms, **Figure 1C**; (Abe et al., 2011; Steel et al., 2016). Hence, in this task, success depended on the ability to reduce the Error by approximating the Target_Force_ as quickly and accurately as possible.

**Figure 1.**
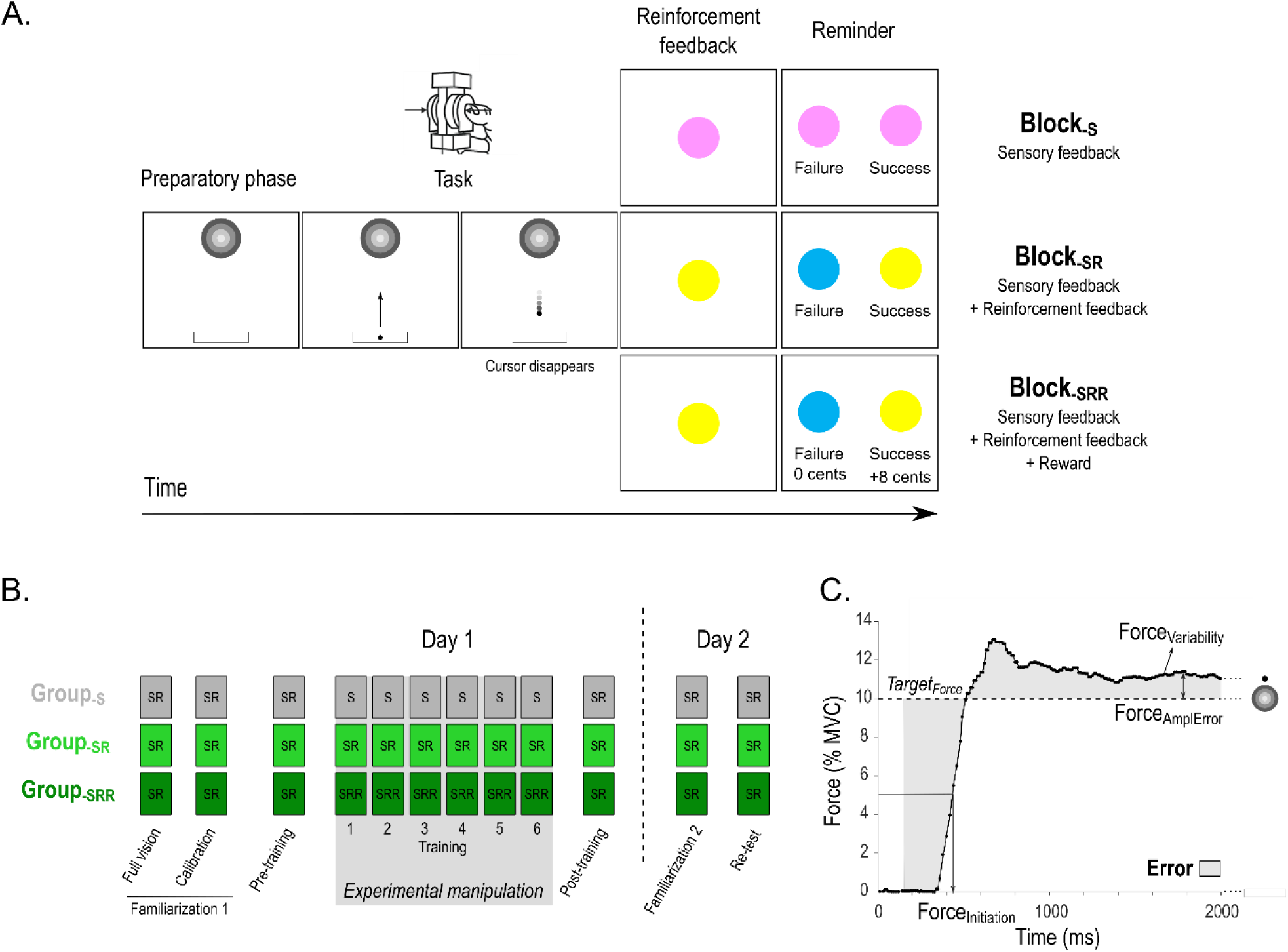
**A) Time course of a trial in the motor skill learning task.** Each trial started with the appearance of a sidebar and a target. After a variable preparatory phase (800-1000ms), a cursor appeared in the sidebar, playing the role of a “Go” signal. At this moment, participants were required to pinch the force transducer to bring the cursor into the target as quickly as possible and maintain it there until the end of the task (2000ms). Notably, on most trials, the cursor disappeared halfway towards the target (as displayed here). Then, a reinforcement feedback was provided in the form of a colored circle for 1000ms and provided knowledge of performance (Block_-SR_, Block_-SRR_) or was non-informative (Block-S). The reinforcement feedback was determined based on the comparison between the Error on the trial and the individual success threshold (computed in the Calibration block, see Materials and Methods). Finally, each trial ended with a reminder of the color/feedback association and potential reward associated to good performance (1500ms). **B) Experimental procedure.** On Day 1, all participants performed two familiarization blocks in a Block_-SR_ condition. The first one involved full vision of the cursor while the second one provided only partial vision and served to calibrate the difficulty of the task on an individual basis (See Materials and Methods). Then, Pre- and Post-training Block_-SR_ assessments were separated by 6 blocks of training in the condition corresponding to each individual group (Block_-S_ for Group_-S_, Block_-SR_ for Group_-SR_ and Block_-SRR_ for Group_-SRR_). Day 2 consisted in a Familiarization block followed by a Re-test assessment (4 Block_-SR_ pooled together). **C) Example of a force profile.** Force applied (in % of MVC) during the task. Participants were asked to approximate the Target_Force_ as quickly and accurately as possible to minimize the Error (grey shaded area). As shown on the figure, this Error depended on the speed of force initiation (Force_Initiation_) and on the accuracy of the maintained force, as reflected by its amplitude with respect to the Target_Force_ (Force_AmplError_) and its variability (Force_Variability_). Note that the first 150ms of each trial were not considered for the computation of the Error.

**Table 1.** Group features and muscle fatigue in the three experimental groups (mean ± SE). The 2 last columns provide the results of one-way ANOVAs’ ran with the factor Group_TYPE_.

|  | **Group-s**<br>**(n=30)** | **Group-SR**<br>**(n=30)** | **Group-SRR**<br>**(n=30)** | **F<sub>(2,87)</sub>** | **p** |
| --- | --- | --- | --- | --- | --- |
| Age<br>(years) | 23.9 ± 0.67 | 23.3 ± 0.50 | 23.9 ± 0.43 | 0.34 | 0.71 |
| Gender<br>(number of females) | 19 | 19 | 20 | / | / |
| Success Threshold<br>(% MVC) | 2.8 ± 0.01 | 2.8 ± 0.01 | 2.9 ± 0.01 | 0.13 | 0.88 |
| Target<sub>Force</sub><br>(Newtons) | 5.3 ± 0.30 | 4.7 ± 0.25 | 5.1 ± 0.21 | 1.16 | 0.31 |
| Sensitivity to reward (score) | 37.1 ± 1.18 | 35.6 ± 1.11 | 37.5 ± 1.14 | 0.79 | 0.46 |
| Sensitivity to punishment (score) | 42.3 ± 1.59 | 42.0 ± 1.59 | 40.9 ± 1.46 | 0.22 | 0.81 |
| Muscle fatigue – Day 1<br>(MVC<sub>POST</sub> in % of MVC<sub>PRE</sub>) | 95.7 ± 2.39 | 99.0 ± 2.61 | 98.6 ± 2.57 | 0.51 | 0.60 |
| Muscle fatigue – Day 2<br>(MVC<sub>POST</sub> in % of MVC<sub>PRE</sub>) | 94.6 ± 2.74 | 93.9 ± 1.52 | 97.0 ± 1.89 | 0.60 | 0.55 |

### Reward improves motor skill learning

Participants’ initial performance was comparable in all groups: the Error in the Pre-training block equaled 3.14 ± 0.18 % MVC in Group_-S_, 3.33 ± 0.17 % MVC in Group_-SR_ and 3.30 ± 0.15 % MVC in Group_-SRR_ (one-way ANOVA: F_(2,87)_ = 0.37, p = 0.69, partial η^2^ = 0.0084; **Figure 2A**). In contrast, skill learning, estimated as the training-related reduction in Error on Day 1 (Post-training expressed in % of Pre-training) varied as a function of the group (**Figure 2B)**. As such, learning was stronger in the Group_-SRR_ compared to the two other groups (ANOVA: F_(2,87)_ = 4.41, p = 0.015, partial η^2^ = 0.092; post-hocs: Group_-SRR_ vs. Group_-SR_: p = 0.014, Cohen’s d = 0.60; Group_-SRR_ vs. Group_-S_: p=0.010, d = 0.98), with no significant difference between Group_-S_ and Group_-SR_ (p = 0.91, d = 0.025). This was confirmed by a subsequent analysis showing that learning was significant in the Group_-SRR_ (Post-training = 80.7 ± 3.5 % of Pre-training; single-sample t-test against 100 %: t_(29)_ = -5.49, p < 0.00001, d = -1.42), but not in Group_-S_ (Post-training = 103.9 ± 5.15 % of Pre-training; t_(29)_ = 0.75, p = 0.46, d = 0.19) or in Group_-SR_ (Post-training = 102.9 ± 8.80 % of Pre-training; t_(29)_ = 0.33, p = 0.745, d = 0.085). Skill maintenance on Day 2, estimated as the Error at Re-test in percentage of Pre-training, was not significantly different between the groups (F_(2,87)_ = 1.96, p = 0.15, partial η^2^ = 0.043; **Figure 2C**). However, in Group_-SRR_, we found that the Error at Re-test remained lower than at Pre-training (Re-test = 85.6 ± 5.01 % of Pre-training; single-sample t-test against 100 %: t_(29)_ = -2.88, p < 0.0073, d = -0.74) demonstrating that the skill was maintained, while this effect was not significant in the two other groups (Group_-S_: Re-test = 100.5 ± 4.63 % of Pre-training; t_(29)_ = 0.11, p = 0.92, d = -0.12, Group_-SR_: Re-test = 97.0± 6.82 % of Pre-training; t_(29)_ = -0.45, p = 0.66, d = -0.028). Hence, while reinforcement alone did not contribute to reduce the Error in this task, its combination with reward successfully helped participants to learn and maintain the skill, as also evident when considering the averaged success rates (**Figure 2D**) and individual force profiles (**Figure 2E**).

**Figure 2.**
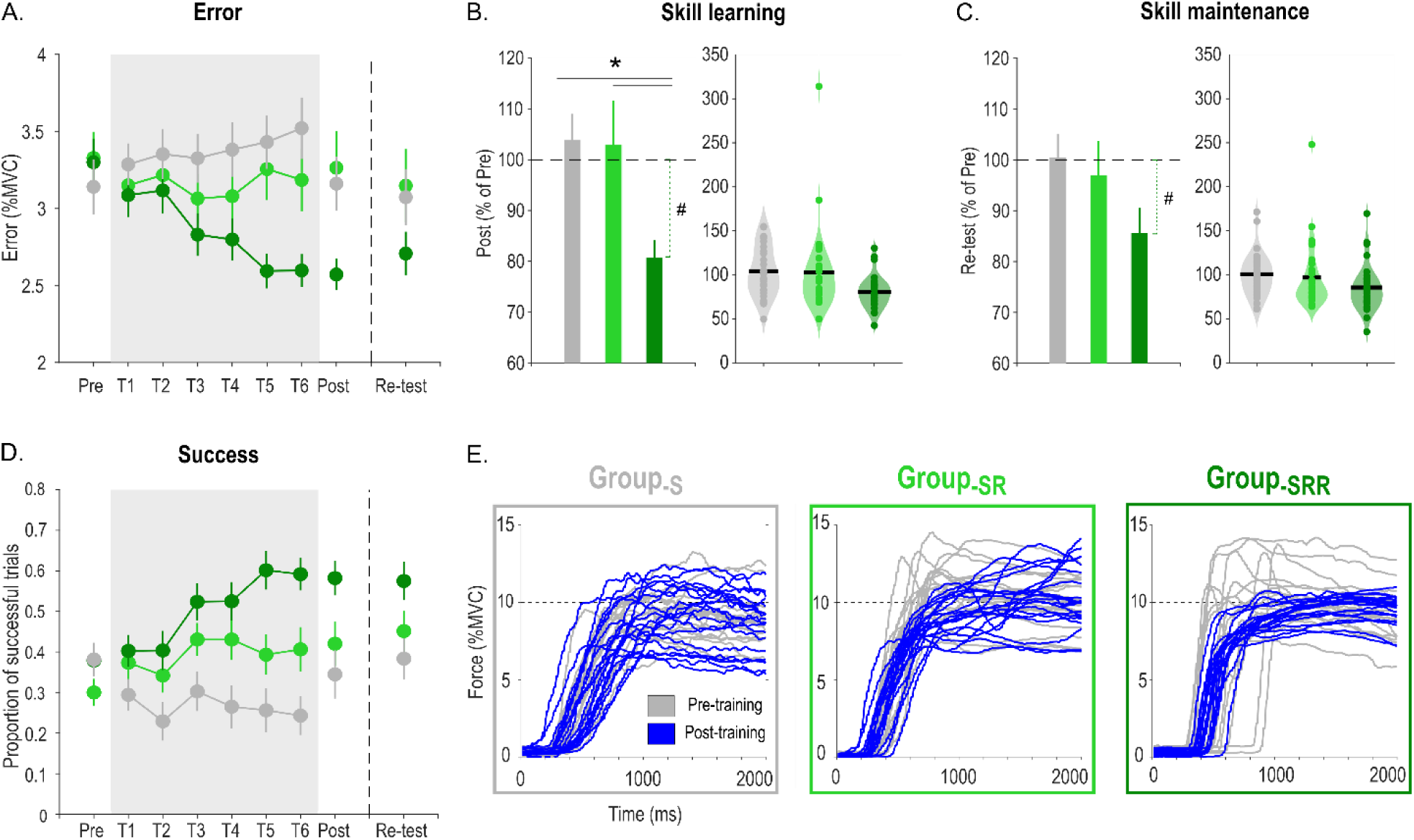
**A) Error.** Average Error is represented across practice for the three experimental groups (grey: Group_-S_, light green: Group_-SR_, dark green: Group_-SRR_). The grey shaded area highlights the blocks concerned by the reinforcement manipulation. The remaining blocks were performed with knowledge of performance only (*i.e.,* in a Block_-SR_ setting). **B) Skill learning.** Bar plot (left) and violin plot (right, each dot = one subject) representing skill learning (quantified as the Error in Post-training blocks expressed in percentage of Pre-training blocks) in the three experimental groups. Skill learning was significantly enhanced in Group_-SRR_ compared to the two other groups. This result remained significant when removing the subject showing an extreme value in the Group_-SR_ (ANOVA: F_(2,86)_ = 6.44, p = 0.0025, partial η^2^ = 0.13; post-hocs; Group-SRR vs. Group-SR: p = 0.027; Group-SRR vs. Group-S: p = 0.00064; Group-SR vs. Group-S: p = 0.21). **C) Skill maintenance.** Bar plot (left) and violin plot (right) representing skill maintenance quantified as the Error in Re-test blocks expressed in percentage of Pre-training blocks) in the three experimental groups. **D) Success.** Proportion of successful trials for each block. **E) Force profiles.** Individual force profiles of one representative subject of Group_-S_ (left), Group_-SR_ (middle) and Group_-SRR_ (right) in the Pre- (grey) and Post-training blocks (blue). *: significant difference between groups (p<0.05). #: significant difference within a group between normalized Post-training Error and a constant value of 100% (p<0.017 to account for multiple comparisons).

### Reward boosts reinforcement-related adjustments during motor skill learning

To identify the mechanisms at the basis of the effect of reward on motor learning, we quantified how much participants adjusted motor commands based on reinforcement or sensory feedback at the single-trial level. This allowed us to estimate how subjects relied on each type of feedback on a trial-by-trial basis and how this behaviour was affected by reward. In order to investigate reinforcement-related adjustments in motor commands, we computed the absolute between-trial change (BTC) in Error (Error_BTC_ = |Error_n+1_-Error_n_|) following successful or failed trial_n_ of similar Error in the three groups (Materials and Methods, see also Pekny et al., 2015 and Uehara et al., 2019 for similar approaches in reaching tasks). Comparing Error_BTC_ following successful and failed trials allowed us to estimate how much participants modified their force profile based on the reinforcement feedback. Notably, considering changes in the Error in absolute terms allowed us to explore the effect of reward on the magnitude of the adjustments in the different groups, regardless of their directionality (increase or decrease in the Error). Interestingly, we found that the magnitude of these adjustments (expressed as Error_BTC_ following a Failure relative to Error_BTC_ following a Success) differed depending on the Group_TYPE_ during Day 1 training (F_(2,84)_ = 10.27, p < 0.001, partial η^2^ = 0.20; **Figure 3A**). As expected, participants of the Group_-SR_ adjusted their force profile depending on the reinforcement feedback, while participants of the Group_-S_ were unable to do so (post-hocs; Group_-S_ vs. Group_-SR_: p = 0.022, d = -0.68). Interestingly, this ability to adjust motor commands based on the reinforcement was amplified by reward (Group_-SR_ vs. Group_-SRR_: p = 0.036, d = -0.57). This result suggests that one mechanism through which reward improves motor learning is the potentiation of reinforcement-related adjustments in motor commands. To confirm this idea, we evaluated the relationship between the magnitude of reinforcement-based changes in motor commands and the average success rate in the following trial across all subjects. Consistently, we found that the magnitude of reinforcement-related adjustments was a strong predictor of the probability of success at the subsequent trial (R^2^ = 0.62; p = 1.5 x 10^-19^; **Figure 3B**), confirming that these adjustments were functionally relevant. Hence, these data suggest that the effect of reward on motor skill learning relies on the ability to adjust movements based on the reinforcement feedback.

**Figure 3.**
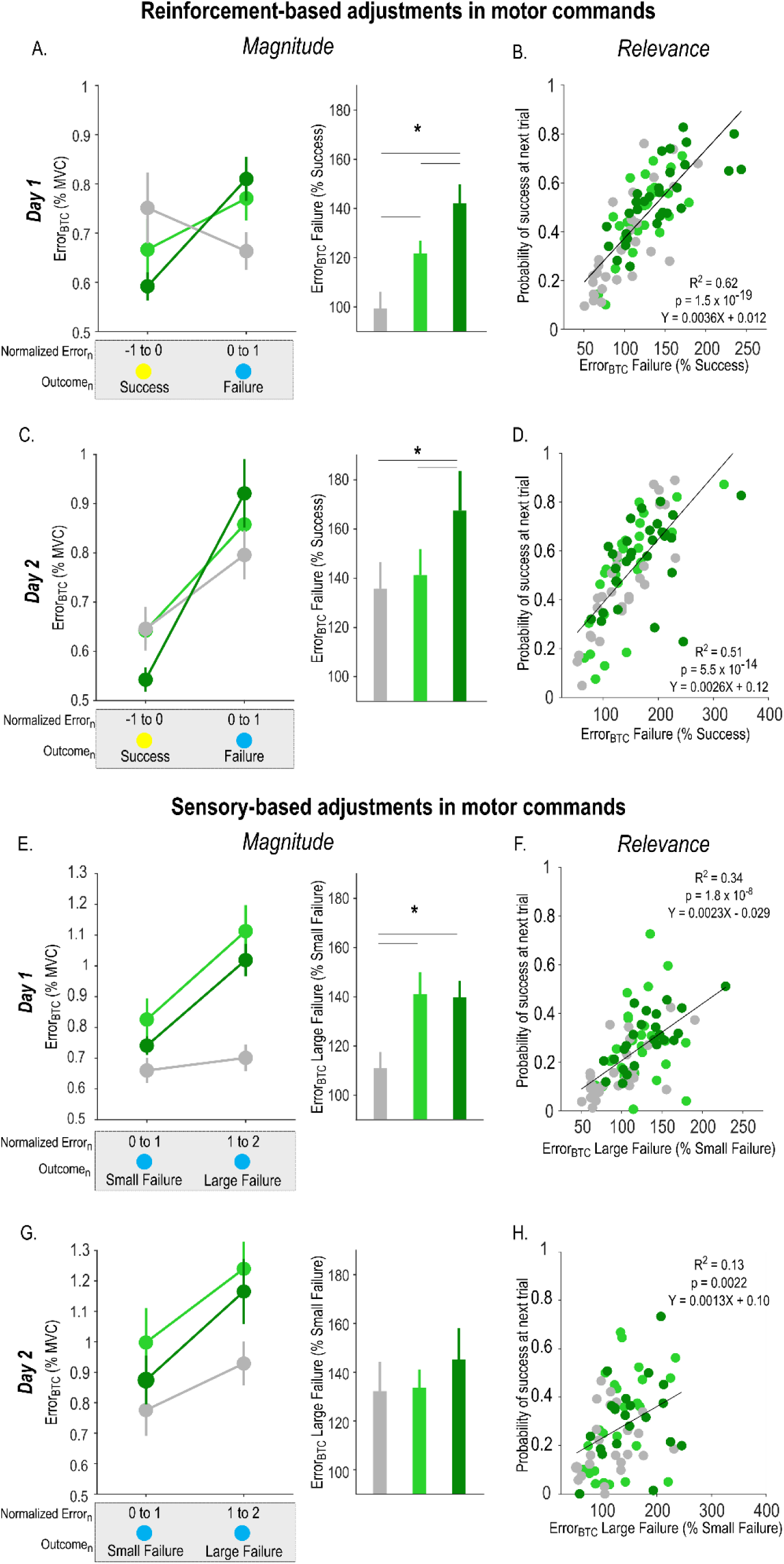
**Between-trial adjustments in the Error. (A)** Reinforcement-based adjustments in the Error during Day 1 training. Absolute between-trial adjustments in the Error (Error_BTC_ = |Error_n+1_-Error_n_|) according to the reinforcement feedback (*i.e.,* Success or Failure) encountered at trial_n_ in the three Group_TYPES_. Notably, these bins of trials where constituted based on the success threshold-normalized Error at trial_n_ in order to compare adjustments in motor commands following trials of similar Error in the three groups (left panel, see Materials and Methods). Reinforcement-based adjustments (Error_BTC_ after Failure in percentage of Error_BTC_ after Success) were compared in the three Group_TYPES_ (right panel). **(B)** Correlations between the magnitude of reinforcement-based adjustments in the Error and the probability of success on the next trial, showing the relevance of these adjustments for task success. Each dot represents a subject. **(C, D)** Same for Day 2 training. Note that reinforcement-based adjustments in motor commands remained amplified in Group_SRR_, despite the absence of reward on Day 2. **(E)** Sensory-based adjustments in the Error during Day 1 training. Error_BTC_ following trials_n_ with Failures of different Error magnitudes (left panel). Sensory-based adjustments (Error_BTC_ after Large Failure in percentage of Error_BTC_ after Small Failure) were compared in the three Group_TYPES_ (right panel). **(F)** Correlations between the magnitude of sensory-based adjustments in the Error and the probability of success on the next trial, showing the relevance of these adjustments for task success. **(G, H)** Same for Day 2 training. *: p < 0.05.

In a second step, we asked whether such single-trial effects were maintained on Day 2, while all participants performed the task with sensory and reinforcement feedback, but in the absence of reward (*i.e.*, in a Block_-SR_ setting). Strikingly, we found that reinforcement-based adjustments in motor commands remained amplified in Group_-SRR_ compared to Group_-S_ and Group_-SR_ (F_(2,78)_ = 3.53, p = 0.034, partial η^2^ = 0.083; post-hocs; Group_-S_ vs. Group_-SRR_: p = 0.017, d = - 0.66, Group_-SR_ vs. Group_-SRR_: p = 0.039, d = -0.56, **Figure 3C**). There was no difference between Group_-S_ and Group_-SR_ (p = 0.72, d = -0.10). Here again, the magnitude of reinforcement-based adjustments correlated with the success in the next trial (R^2^ = 0.51; p=5.5 x 10^-14^; **Figure 3D**). Hence, the effect of reward on reinforcement-based adjustments can persist on a subsequent session of training, even after reward removal.

Then, we evaluated how participants adjusted their movements based on sensory feedback. We reasoned that the extent to which participants relied on the somatosensory feedback to adjust their movements could be estimated by computing Error_BTC_ following failed trials of different Error magnitudes (*i.e.,* small or large Failure). In other words, we contrasted Error_BTC_ following trials with the same reinforcement feedback (*i.e.,* Failure) but with different somatosensory experiences. Interestingly, we found a Group_TYPE_ effect (F_(2,76)_ = 5.05, p = 0.0087, partial η^2^ = 0.12; **Figure 3E)** that was driven by differences between Group_-S_ and the two other groups (post-hocs; Group_-S_ vs. Group_-SR_: p = 0.0056, d = -0.73, Group_-S_ vs. Group_-SRR_: p = 0.011, d = -0.85). This indicates that while subjects of the Group_-SR_ and Group_-SRR_ adjusted the Error differently depending on the sensory feedback, participants of the Group_-S_ were less able to do so, suggesting that training with reinforcement feedback allowed participants to be more sensitive to the sensory feedback, as previously suggested (Bernardi et al., 2015; Galea et al., 2015). Importantly though, we did not find any difference between Group_-SR_ and Group_-SRR_ (p = 0.90, d = -0.033), suggesting that reward did not modify these sensory-based adjustments in motor commands. Then, similarly as for the reinforcement-based changes, we found that the magnitude of sensory-based adjustments correlated with the subsequent probability of success (R^2^ = 0.34, p = 1.8 x 10^8^; **Figure 3F**), demonstrating that these adjustments were also relevant in the learning process. Notably, on Day 2, sensory-based adjustments were not different between the Group_TYPES_ (F_(2,68)_ = 0.41, p = 0.67, partial η^2^ = 0.012, **Figure 3G**), but still significantly correlated with the probability of success (R^2^ = 0.13, p = 0.0022; **Figure 3H**). Thus, our data suggest that despite the relevance of the somatosensory feedback in the present task, reward did not modulate the extent to which participants relied on it to improve their movements. Rather, it appears that reward caused a specific and persistent potentiation of reinforcement-based adjustments in motor commands.

### Reward boosts reinforcement-based adjustments at a specific level of motor control

As a last step, we asked whether the effect of reward on between-trial adjustments in motor commands concerned all aspects of force control, or only some specific motor components. To do so, we investigated how reinforcement and sensory feedback shaped adjustments in the speed and accuracy of force production in the three Group_TYPES_ by dissecting each force profile into three separate components (**Figure 1B**). To evaluate the speed at which the force was generated, we computed the time required for force initiation (*i.e.*, the time required to reach half of the Target_Force_: Force_Initiation_). To assess the accuracy of the force, we computed the force difference between the average amplitude of the generated force and the Target_Force_ (Force_AmplError_), and the variability (standard deviation/mean) of the maintained force (Force_Variability_). Notably, both indicators of force accuracy were computed in the second half of the trial (*i.e.*, the last 1000 ms), well after force initiation, when participants maintained a stable level of force.

We compared between-trial changes in Force_Initiation_ (Force_Initiation-BTC_), Force_AmplError_ (Force_AmplError-BTC_), and Force_Variability_ (Force_Variability-BTC_) following Success or Failure trials of similar Error magnitude in the three groups. The presence of the reinforcement feedback impacted the modulation of initiation speed (expressed as Force_Initiation-BTC_ following a Failure in percentage of Force_Initiation-BTC_ following a Success; F_(2,84)_ = 8.50, p < 0.001, partial η^2^ = 0.17; post-hocs: Group_-S_ vs. Group_-SR_: p = 0.0011, d = -0.84, Group_-S_ vs. Group_-SRR_: p < 0.001, d = -1.09; **Figure 4A**). Interestingly though, we did not find any effect of reward on the reinforcement-based adjustment of speed (Group_-SR_ vs. Group_-SRR_: p = 0.78, d = -0.072). However, there were significant effects of reward on reinforcement-related changes in Force_AmplError-BTC_ (F_(2,84)_ = 9.54, p < 0.001, partial η^2^ = 0.19; **Figure 4B**). As such, participants of the Group_-SRR_ modulated more the Force_AmplError_ according to the reinforcement feedback than subjects of the two other groups (Group_-S_ vs. Group_-SRR_: p < 0.001, d = -1.04, Group_-SR_ vs. Group_-SRR_: p = 0.018, d = -0.70). Notably, there was also a trend for Group_-SR_ to be different from Group_-S_ (p = 0.064, d = -0.50). Finally, we found no Group_TYPE_ effect on the Force_Variability-BTC_ (F_(2,84)_ = 0.81, p = 0.45, partial η^2^ = 0.020; **Figure 4C**). Hence, while reward strongly influenced the reinforcement-based adjustments of force amplitude, it did not modulate the between-trial regulation of the speed at which the force was initiated as well as the variability of the maintained force. This suggests that the effect of reward on reinforcement-related adjustments was not global (*i.e.*, affecting all aspects of the movement) but rather specific to force amplitude.

**Figure 4.**
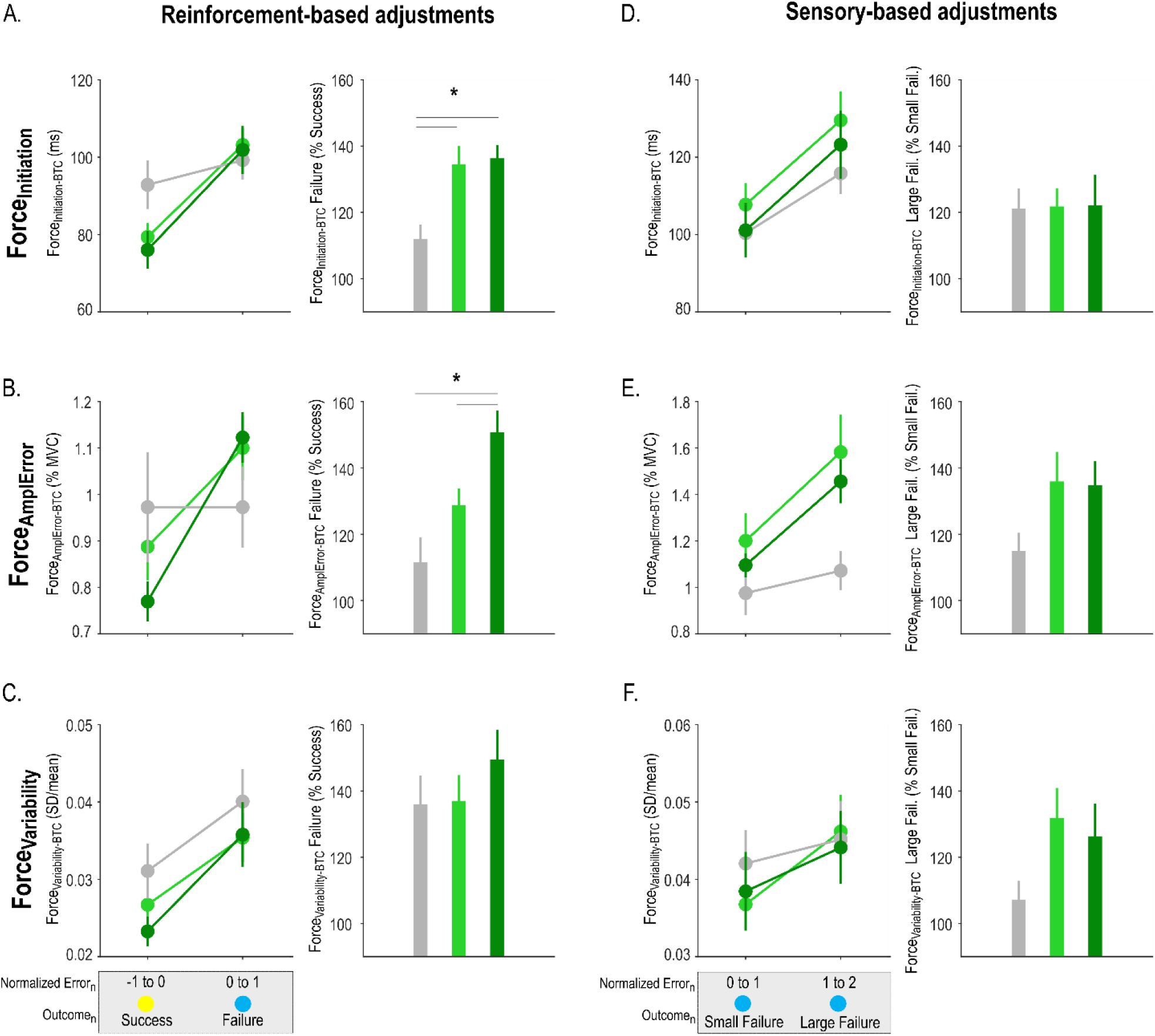
**Between-trial adjustments in initiation time, amplitude error and variability. Reinforcement-based adjustments in the Force_Initiation_ (A), Force_AmplError_ (B) and Force_Variability_ (C).** Absolute between-trial changes (BTC) for each motor component (Force_BTC_ = |Force_n+1_-Force_n_|) according to the reinforcement feedback (*i.e.,* Success or Failure) encountered at trial_n_ in the three Group_TYPES_. Notably, these bins of trials where constituted based on the success threshold-normalized Error at trial_n_ (left panel). Reinforcement-based adjustments (Force_BTC_ after Failure in percentage of Force_BTC_ after Success) in the three Group_TYPES_ (right panel). **Sensory-based adjustments in the Force_Initiation_ (D), Force_AmplError_ (E) and Force_Variability_ (F).** Force_BTC_ following trials_n_ with Failures of different Error magnitudes (left panel). Sensory-based adjustments (Force_BTC_ after Large Failure in percentage of Force_BTC_ after Small Failure) in the three Group_TYPES_ (right panel). *: p < 0.05.

We also considered the effect of the sensory feedback on between-trial adjustments by comparing Force_Initiation-BTC_, Force_AmplError-BTC_, and Force_Variability-BTC_ following failed trials of different Error magnitudes (*i.e.,* small or large Failure). Here, we did not find any significant difference in the way participants adjusted individual motor components in response to the sensory feedback (**Figure 4D, 4E, 4F**; Force_Initiation-BTC_: F_(2,76)_ = 0.10, p = 0.90, partial η^2^ = 0.0026; Force_AmplError-BTC_: F_(2,76)_ = 2.57, p = 0.083, partial η^2^ = 0.063; Force_Variability-BTC_: Kruskal-Wallis ANOVA, H_(2)_=5.73, p = 0.057, partial η^2^ = 0.061). This analysis supports the idea that reward did not increase the sensitivity to the sensory feedback, but rather boosted specific adjustments in motor commands in response to the reinforcement feedback.

Finally, as a control analysis, we characterized the respective influence of each motor component in the Error, which determined task success. As such, in addition to representing different levels of force control (*i.e.,* initiation, amplitude and variability), the motor components evaluated here may also bear different relevance for task success (van der Kooij et al., 2021). For each participant, we ran separate partial linear regressions on the Error data with Force_Initiation_, Force_AmplError_ or Force_Variability_ as predictors, while controlling for the effect of the other motor components. Interestingly, we found that Force_AmplError_ explained the largest part of variance in the Error (r = 0.96 ± 0.003; p<0.05 in 90/90 subjects). Force_Initiation_ also explained a large part of variance in the Error (r = 0.81 ± 0.01; p<0.05 in 90/90 subjects), while Force_Variability_ explained a smaller, yet significant in most subjects, part of variance (r = 0.22 ± 0.03; p<0.05 in 68/90 subjects). Hence, although all motor parameters were relevant for task success, the Force_AmplError_ was the most influential factor.

Altogether, our results demonstrate that reward potentiates reinforcement-based adjustments in motor commands and that this effect persists even after reward removal, on the subsequent day. The data also show that this effect does not concern all components of the movement but specifically the amplitude of the force which was the most relevant factor for task success.

## 3. Discussion

In this study, we investigated the net effect of reward on motor learning while controlling for the reinforcement feedback received by the participants. Our results provide evidence that reward can improve motor skill learning and that this effect is related to a specific potentiation of reinforcement-related adjustments in motor commands. Strikingly, the potentiation of such adjustments persisted on a subsequent day in the absence of reward. Moreover, such boosting of reinforcement-based adjustments did not concern all components of force production but only the amplitude, which was the most relevant one for task success. These findings shed light on the mechanisms through which reward can durably enhance motor performance. They also lay the groundwork for future rehabilitation strategies involving optimized sensory and reinforcement feedbacks.

A main goal of the present study was to explore the net effect of reward on motor skill learning by experimentally dissociating it from the reinforcement feedback. As such, previous motor learning studies have often coupled reinforcement and reward (*e.g.,* Abe et al., 2011; Steel et al., 2019, 2016; Wachter et al., 2009; Wilkinson et al., 2015), based on the underlying assumption that receiving knowledge of performance (*e.g.,* points or binary feedback) provides a form of intrinsic reward that can by itself increase motivation to perform well (Leow et al., 2018). However, in addition to providing some form of intrinsic reward, reinforcement feedback also provides a learning signal to the motor system, that can influence motor learning (Bernardi et al., 2015; Galea et al., 2015; Huang et al., 2011; Kim et al., 2019; Leow et al., 2018; Mawase et al., 2017; Nikooyan et al., 2015; Shmuelof et al., 2012; Therrien et al., 2016; Uehara et al., 2018). In order to assess the net impact of motivation on motor skill learning, we therefore compared groups of participants trained with different monetary rewards but with the exact same reinforcement feedback. We found that motivation by reward allowed marked improvements in motor performance that were maintained after reward removal and even 24 hours later (**Figure 2**). Notably, this was the case despite the fact that reinforcement alone was not sufficient to influence motor learning in our task. This demonstrates that the motivational context experienced during training can by itself strongly impact motor skill learning, beyond knowledge of performance.

The prospect of obtaining rewards for good performance enhances motivation but does not provide any additional learning signal to the motor system (Berke, 2018). Yet, it may boost the reliance on sensory and/or reinforcement feedbacks (Kim et al., 2019). To explore this possibility, we developed an analysis allowing us to investigate how participants adjusted their motor commands based on sensory or reinforcement feedbacks while controlling for differences in performance between the groups (see Materials and Methods for more details). Interestingly, we found that reward specifically boosted reinforcement-based adjustments, while sensory-based adjustments remained unaffected by reward (Figure 3). This was the case despite the fact that both types of feedback were relevant to improve motor performance at the single-trial level (**Figure 3B, D, F, H**). This result suggests that reward increases the reliance on reinforcement information during the learning process, with little effect on sensory-based adjustments. Interestingly, this finding may explain why tasks that strongly emphasize sensory-based learning (over reinforcement-based learning; (Cashaback et al., 2017; Izawa and Shadmehr, 2011), often show less sensitivity to motivation. Accordingly, monetary reward shows little impact on sensorimotor adaptation (Galea et al., 2015; Hill et al., 2020) and on motor skill acquisition in tasks that strongly rely on sensory feedback (*e.g.,* Abe et al., 2011; Steel et al., 2016; Widmer et al., 2016). More generally, our results support the idea that one mechanism through which motivation can improve motor learning is the specific potentiation of reinforcement-based adjustments in motor commands.

The finding of a reward-dependent boosting of reinforcement-based adjustments is in line with previous neuroimaging results showing that reward increases reinforcement-related activity in the striatum in the context of motor learning (Widmer et al., 2016). This reward-driven increase in striatal activity is reduced after a stroke (even when the striatum is unlesioned), a process that may contribute to the motor learning deficits observed in these patients (Widmer et al., 2019). Moreover, such reward-dependent modulation of motor adjustments has been shown to rely on dopamine (Galea et al., 2013; Pekny et al., 2015), a key neurotransmitter of the striatal circuitry. Based on these elements and on the causal role of the striatum in reinforcement-based adjustments in motor commands (Nakamura and Hikosaka, 2006; Williams and Eskandar, 2006), we suspect that this region may be crucial for the beneficial effect of reward observed in the present study. Notably, the cerebellum (Carta et al., 2019; Heffley et al., 2018; Sendhilnathan et al., 2020; Vassiliadis et al., 2019; Wagner et al., 2017) and frontal areas (Dayan et al., 2018, 2014; Hamel et al., 2018; Palidis et al., 2019; Ramakrishnan et al., 2017; Sidarta et al., 2016) are also likely to contribute to reward-based motor learning. Further investigations are required to better delineate the neurophysiological bases of reward-related improvements in motor learning.

The beneficial effect of reward on single-trial adjustments was maintained on day 2, even after reward removal. As in day 1 training, reinforcement-based adjustments were boosted while sensory-based adjustments remained unchanged by reward. This persistent change in the specific reaction to the reinforcement feedback after reward removal is suggestive of an associative learning process. In associative learning, presentation of a neutral stimulus (*i.e.*, a conditioned stimulus) that has been consistently paired with a rewarding stimulus (*i.e.*, an unconditioned stimulus) during a training period elicits a behaviour that was initially only generated in reaction to the reward (Pavlov, 1927; Rescorla and Wagner, 1972). Following this framework, it is possible that the repetitive pairing of the reinforcement feedback with the reward during training induced an implicit association between the two events that remained evident when the reward was removed. This could explain why strong reinforcement-specific adjustments were maintained on day 2 in the reward group, even though no rewards were at stake anymore. Such associative learning processes are known to strongly influence autonomic responses (Pool et al., 2019), inhibitory control (Avraham et al., 2020b; Lindström et al., 2019; Verbruggen et al., 2014), decision making (Lindstrom et al., 2019) and even sensorimotor adaptation (Avraham et al., 2020) in humans. We propose that associative learning may also contribute to the durable influence of motivation on motor skill learning (Abe et al., 2011; Sporn et al., 2020).

In order to better characterize the effect of reward on motor learning, we considered separately the different components of the movement and found that force amplitude was the most strongly affected, while the speed of initiation and force variability remained largely insensitive to reward. This suggests that reward can have a selective influence on the regulation of a specific component of motor control. Importantly, an estimation of the respective influence of each motor component on task success also showed that force amplitude was the most relevant component for the task. Notably, the specificity of the effect of reward on the regulation of one motor component is in accordance with the idea that multidimensional motor tasks (*i.e.,* requiring the control of multiple motor components) can be decomposed in subtasks that are learned separately in the motor system (Ghahramani and Wolpert, 1997) In this framework, learning of the different motor components may depend on their respective relevance for task success (Ghahramani and Wolpert, 1997; van der Kooij et al., 2021). Such task relevance may be estimated based on a priori knowledge of the task (*e.g.,* following instructions; Popp et al., 2020) and through the reliance on a credit assignment system allowing to estimate the particular influence of each motor component on task success through trial and error (McDougle et al., 2016; Parvin et al., 2018). Based on this idea, we believe that the strong relationship between the amplitude of the force and task success in the present task pushed participants of the reward group to largely modulate this component based on the reinforcement feedback. If this is the case, this would suggest that it is possible to affect the training of specific motor abilities by modulating the weight of individual motor components in the computation of the reinforcement feedback, an aspect that could be exploited in future rehabilitation protocols. Alternatively, reward might have specifically modulated the amplitude of the force, independently of the relevance of this parameter. Although the present study cannot rule out this hypothesis, we believe that such interpretation is unlikely given previous demonstration that reward can improve several aspects of motor control concomitantly (Codol et al., 2020; Manohar et al., 2015). Another possibility is that reinforcement feedback alone was sufficient to maximally modulate initiation time and variability in this task, precluding us from observing a difference with the reward-based training because of some form of ceiling effect. Further studies are required to disentangle these potentially co-existing interpretations to guide the development of component-specific rehabilitation therapies (Norman et al., 2017).

## 4. Materials and Methods

### 4.1. Participants

A total of ninety right-handed healthy volunteers participated in the present study (58 women, 23.7 ± 0.3 years old; mean ± SE). Handedness was determined via a shortened version of the Edinburgh Handedness inventory Oldfield (Oldfield, 1971). None of the participants suffered from any neurological or psychiatric disorder, nor were they taking any centrally-acting medication. All participants gave their written informed consent in accordance with the Ethics Committee of the Université Catholique de Louvain (approval number: 2018/22MAI/219) and the principles of the Declaration of Helsinki. Subjects were financially compensated for their participation. Finally, all participants were asked to fill out a French adaptation of the Sensitivity to Punishment and Sensitivity to Reward Questionnaire (SPSRQ; Lardi et al., 2008; Torrubia et al., 2001).

### 4.2 Motor skill learning task

#### 4.2.1 General aspects

Participants were seated approximately 60 cm in front of a computer screen (refresh rate = 100 Hz) with their right forearm positioned at a right angle of the table. The task was developed on Matlab 7.5 (the Mathworks, Natick, Massachusetts, USA) exploiting the Psychophysics Toolbox extensions (Brainard, 1997; Pelli, 1997) and consisted in an adaptation of previously used motor learning tasks (Abe et al., 2011; Mawase et al., 2017; Steel et al., 2016). The task required participants to squeeze a force transducer (Arsalis, Belgium) between the index and the thumb to control the one-dimension motion of a cursor displayed on the screen. Increasing force resulted in the cursor moving vertically and upward. Each trial started with a preparatory phase in which a sidebar appeared at the bottom of the screen and a target at the top (**Figure 1A**). After a variable time interval, a cursor popped up in the sidebar and participants had to pinch the transducer to move the cursor as quickly as possible from the sidebar to the target and maintain it there for the rest of the task. The level of force required to reach the target (Target_Force_) was individualized for each participant and set at 10% of maximum voluntary contraction (MVC). Notably, squeezing the transducer before the appearance of the cursor was considered as an anticipation and therefore led to an interruption of the trial. Such trials were discarded from further analyses. At the end of each trial, a binary reinforcement feedback represented by a colored circle was provided to the subject followed by a reminder of the color/feedback association and potential monetary reward associated to good performance (see Reinforcement feedback section below).

#### 4.2.2. Sensory feedback

We provided only limited visual feedback to the participants (Mawase et al., 2017). As such, on most trials (90%), the cursor disappeared shortly after the subject started to squeeze the force transducer (partial vision trials): it became invisible as soon as the generated force became larger than half of the Target_Force_ (*i.e.,* 5% of MVC). Conversely, the remaining trials (10%) provided a continuous vision of the cursor (full vision trials). Therefore, on most trials, participants had limited visual information and had to rely exclusively on somatosensory feedback to generate the Target_Force_. Importantly, full vision trials were not considered in the analyses.

#### 4.2.3 Reinforcement feedback

At the end of each trial, subjects were presented with a binary reinforcement feedback indicating performance. Success on the task was determined based on the Error; that is, the absolute force difference between the Target_Force_ and the exerted force (**Figure 1B**; Abe et al., 2011; Steel et al., 2016). The Error was computed for each frame refresh (*i.e.,* at 100Hz) from 150 ms to the end of the trial and then averaged for each trial (Steel et al., 2016) and expressed in percentage of MVC. This indicator of performance allowed us to classify a trial as successful or not based on an individualized success threshold (see below). When the Error on a given trial was below the threshold, the trial was considered as successful, and when it was above the threshold, the trial was considered as failed. Hence, task success depended on the ability to reduce the Error by approximating the Target_Force_ as quickly and accurately as possible. Importantly, participants were told explicitly that both speed and accuracy were taken into account to determine task success. In summary, to be successful, participants knew that they had to quickly initiate the force and be as accurate as possible in reproducing the Target_Force._

In different blocks of trials, we manipulated the reinforcement feedback and reward provided during training. In Block_-S_, the reinforcement feedback was non-informative (magenta circle regardless of performance), and participants could only rely on somatosensory feedback to perform the task. In Block_-SR_, the reinforcement feedback consisted in a yellow (representing Success) or blue circle (representing Failure), providing knowledge of performance (**Figure 1A**). In Block_-SRR_, this knowledge of performance was associated to a monetary reward (+8 cents or 0 cent for Success or Failure, respectively). Therefore, contrarily to Block_-S_, Block_-SR_ and Block_-SRR_ provided knowledge of performance and this feedback was associated to a monetary reward in Block_-SRR_.

### 4.3. Experimental procedure

Subjects’ performance was tested for two consecutive days (Day 1 and Day 2; **Figure 1C**). On Day 1, we first measured the individual MVC to calculate the Target_Force_. Notably, MVC was measured before and after both sessions to assess potential muscle fatigue related to the training (see 4.4.3). Participants then performed 2 blocks of Familiarization. In a first block, participants performed 20 full vision trials; it served to familiarize the subjects with the task in a Block_-SR_ setting (Full vision block). Subsequently, all blocks were composed of a mixture of full vision trials (10 % of total trials) and partial vision trials (90 % of total trials). The second Familiarization block consisted in 20 trials and allowed us to determine baseline performance to individualize the difficulty of the task for the rest of the experiment (Calibration block). For every subject, each partial vision trial of the Calibration block was classified in terms of Error from the lowest to the greatest in percentage of MVC. We took the 35^th^ percentile of the Error to determine the individual success threshold. Success thresholds were constrained between 2 and 3.5 % of MVC by asking participants to repeat the Calibration block when the computed threshold was outside these boundaries. Those parameters were determined based on pilot data to obtain coherent learning curves among individuals.

After the Familiarization and Calibration blocks, the first experimental session consisted in 280 trials divided in 8 blocks. All subjects started with a Block_-SR_ of 20 trials to evaluate the performance at Pre-training and similarly ended the session with a Post-training assessment of 20 trials. In between, 6 Training blocks of 40 trials were performed by the participants (**Figure 1B**). During this Training period, individuals were split into 3 separate groups (Group_TYPE_: Group_-S_, Group_-SR_ or Group_-SRR_) depending on the type of blocks they performed during training. As such, Group_-S_ completed Block_-S_, Group_-SR_ performed Block_-SR_ and Group_-SRR_ trained under Block_-SRR_ condition. Contrasting performance in the Pre- and Post-training blocks allowed us to evaluate learning of the skill under the three training conditions. 24 hours later, subjects performed the task again with the same Target_Force_ and success threshold. After a 20 trials Familiarization, participants performed 140 trials split in 4 blocks; all were performed in a Block_-SR_ setting. This Re-test session allowed us to assess skill maintenance 24h after training.

### 4.4. Data and statistical analysis

Statistical analyses were carried out with Matlab 2018a (the Mathworks, Natick, Massachusetts, USA) and Statistica 10 (StatSoft Inc., Tulsa, Oklahoma, USA). In the case of Analysis of Variance (ANOVA), assumption of homogeneity of variance was systematically verified by means of Levene’s test and non-parametric Kruskal-Wallis ANOVA was used when this assumption was violated. Post-hoc comparisons were always conducted using the Fisher’s LSD procedure. The significance level was set at p ≤ 0.05, except in the case of correction for multiple comparisons (see below).

#### 4.4.1. Motor skill learning and maintenance

The main aim of the present study was to evaluate the effect of reward on motor skill learning and maintenance. To assess skill learning, we expressed the median Error at Post-training in percentage of the value obtained at Pre-training. To evaluate skill maintenance, we expressed the median Error during the Re-test session in percentage of Pre-training. First, compared skill learning and maintenance between the groups through one-way ANOVAs with the factor Group_TYPE_. Then, we also explored the significance of skill learning and maintenance within each group by conducting Bonferroni-corrected single sample t-tests on these percentage data against a constant value of 100% (*i.e.,* corresponding to the Pre-training level).

As explained above, task performance depended on both the speed and the accuracy of the produced force (**Figure 1B**). We characterized the effect of reward on these different levels of force control, by evaluating separately the speed of force initiation and the accuracy of the maintained force. To evaluate the speed of force initiation, we measured the force initiation time (Force_Initiation_) which was defined as the delay between the appearance of the cursor and the moment where the applied force reached 5% of MVC (*i.e.,* corresponding to half of the Target_Force_). Force accuracy was evaluated in the second half of the trial (*i.e.,* the last 1000 ms), through two different parameters. First, we computed the Amplitude Error of the force (Force_AmplError_), defined as the absolute difference between the mean force exerted in the last 1000 ms of the trial and the Target_Force_. It reflected how much the amplitude of the maintained force differed from the Target_Force_. Second, force accuracy was also characterized by considering the variability of the maintained force, with high levels of variability causing increases in the Error. To assess force variability (Force_Variability_), we computed the coefficient of variation of the force in the second half of the trial (*i.e.,* standard deviation of force/mean force). In summary, to be successful, participants had to quickly initiate the force (*i.e.,* low Force_Initiation_) and be as accurate as possible (*i.e.,* low Force_AmplError_ and Force_Variability_).

As a control, we verified that the three motor components described above (*i.e.,* Force_Initiation_, Force_AmplError_ and Force_Variability_) were closely related to the Error, and therefore were relevant for task success. To do so, we ran partial linear regressions on the Error data with Force_Initiation_, Force_AmplError_ and Force_Variability_ as predictors to estimate the respective influence of each motor component on the Error, while controlling for the effect of the other components. Interestingly, we found that Force_AmplError_ explained the largest part of variance in the Error (r = 0.96 ± 0.003; p<0.05 in 90/90 subjects). Force_Initiation_ also explained a large part of variance in the Error (r = 0.81 ± 0.01; p<0.05 in 90/90 subjects), while Force_Variability_ explained a smaller, yet significant in most subjects, part of variance (r = 0.22 ± 0.03; p<0.05 in 68/90 subjects). Hence, although all motor parameters were relevant for task success, the Force_AmplError_ was the most influential factor.

#### 4.4.2. Between-trial adjustments in motor commands

A second goal of the present study was to assess the effect of reward on between-trial adjustments in motor commands. Specifically, we aimed at evaluating how motor commands were adjusted based on reinforcement and sensory feedback in our three experimental groups.

To do so, for each trial_n_ we computed the absolute between-trial change (BTC) in Error (Error_BTC_; see Pekny et al., 2015; Uehara et al., 2019 for similar approaches in reaching tasks).

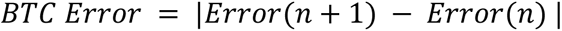

In order to study how much motor commands were adjusted based on previous experience, we compared adjustments in motor commands following trials of different Error magnitudes. To do so, we first subtracted each subject’s individual success threshold to the Error data. Hence, normalized Errors below 0 corresponded to successful trials and normalized Errors above 0 corresponded to failed trials. Then, we split the Error data in consecutive bins of 1 % of MVC and averaged the corresponding Error_BTC_. This allowed us to compare Error_BTC_ following trials of similar Error_n_ across the groups.

As a first step, to better understand how motor commands were adjusted based on the reinforcement feedback, we compared Error_BTC_ following bins of Success or Failure trials of neighboring Error magnitudes (Bin_Success_: -1% < Error_n_ < 0% MVC; Bin_Failure_: 0% < Error_n_ < 1% MVC). Fixing the boundaries of Bin_Success_ and Bin_Failure_ allowed us to compare reinforcement-related adjustments between the groups while controlling for the magnitude of Error_n_; an aspect that might directly influence between-trial adjustments. We then computed reinforcement-based adjustments as the percentage change in Error_BTC_ in Bin_Failure_ compared to Bin_Success_. This index allowed us to determine in a single measure how participants from the different groups adjusted their behavior based on the reinforcement obtained in the previous trial.

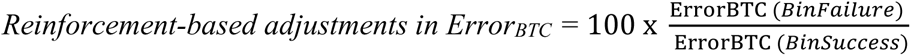

This analysis was conducted separately on the Day 1 and Day 2 data. We had to exclude 3 and 9 participants for Day 1 and Day 2 analyses, respectively, because they had less than 7 trials in at least one of the two bins (remaining subjects on Day 1: Group_-S_ = 29; Group_-SR_ = 28; Group_SRR_ = 30; Day 2: Group_-S_ = 26; Group_-SR_ = 27; Group_-SRR_ = 28). Reinforcement-based changes in Error_BTC_ were compared between the groups through one-way ANOVAs with the factor Group_TYPE_.

As a second step, we evaluated how participants adjusted movements when they could only rely on the sensory feedback. We compared Error_BTC_ following bins of Failure trials of different Error magnitudes (Bin_Small-Failure_: 0% < Error_n_ < 1% MVC; Bin_Large-Failure_: 1% < Error_n_ < 2% MVC). In this case, the reinforcement feedback was the same in the two bins and the only difference between the trials consisted in the magnitude of the Error experienced at trial_n_. This index allowed us to determine how participants adjusted their behavior based on the previous somatosensory experience, in the absence of any difference in the reinforcement feedback obtained.

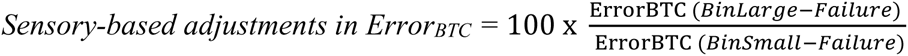

This analysis was first run on the Day 1 data. We had to exclude 12 participants because they had less than 7 trials in at least one of the two bins (remaining subjects: Group_-S_ = 27; Group_-SR_ = 28; Group_-SRR_ = 24). For Day 2, applying the same procedure led to the exclusion of 29 subjects with a lower number of participants in the Group_-SRR_ (15 subjects). For this reason, we ran another analysis where we exceptionally excluded participants only if they had less than 5 trials in one bin. This allowed us to keep a reasonable number of participants in each group (19 subjects excluded; remaining subjects: Group_-S_ = 24; Group_-SR_ = 26; Group_-SRR_ = 21). Notably, both analyses (*i.e.,* with 7-trials or 5-trials cutoff) gave similar results and we only present the latter in the Results section. Sensory-based changes in Error_BTC_ were compared between the groups through a one-way ANOVA with the factor Group_TYPE_.

As a last step, we asked whether the effect of reward on between-trial adjustments in motor commands concerned all aspects of force control, or only specific motor components. To do so, we investigated reinforcement-based and sensory-based adjustments in Force_Initiation_, Force_AmplError_ and Force_Variability_, using the same method described above for the average Error. To assess reinforcement-based adjustments, we contrasted between-trial changes in Force_Initiation_ (Force_Initiation-BTC_), Force_AmplError_ (Force_AmplError-BTC_) and in Force_Variability_ (Force_Variability-BTC_) following Bin_Success_ and Bin_Failure_. Sensory-based adjustments were computed by contrasting Force_Initiation-BTC_, Force_AmplError-BTC_ and Force_Variability-BTC_ following Bin_Small-Failure_ and Bin_Large-Failure_. These data were compared between the groups through one-way ANOVAs with the factor Group_TYPE_.

#### 4.4.3 Group features, muscle fatigue and monetary gains

As a control, we verified that our 3 groups were comparable in terms of age, success threshold, Target_Force_ and Sensitivity to Reward and to Punishment (*i.e.,* as assessed by the SPSRQ questionnaire). As displayed in **Table 1**, one-way ANOVAs on these data did not reveal any significant differences between the groups.

We also assessed muscle fatigue on Day 1 and Day 2 (Derosiere & Perrey 2012, Derosiere et al., 2014) by expressing the MVC obtained after each session (MVC_POST_) in percentage of the MVC measured initially (MVC_PRE_). The relative change of MVC was not different according to the Group_TYPE_ (Day 1, F_(2,87)_ = 0.51, p = 0.60; Day 2, F_(2,87)_ = 0.60, p = 0.55; **Table 1**). As an additional safety check, we wanted to make sure that the decrements in MVC caused by the training period of Day 1 could not impair performance. To test this, we compared MVC_POST_ (expressed in % of MVC_PRE_) with a fixed value of 10% of MVC_PRE_ (*i.e.,* corresponding to the Target_Force_) through Bonferroni-corrected single sample t-tests. This analysis revealed that MVC_POST_ levels were always significantly above the Target_Force_ (Group_-S_: t_(29)_ = 35.84, p < 0.001; Group_-SR_: t_(29)_ = 34.14, p < 0.001 and Group_-SRR_: t_(29)_ = 34.44, p < 0.001). Hence, force decrements caused by the training were comparable between groups and are unlikely to have limited task performance.

In a final step, we checked that the monetary gains obtained at the end of the experiment were similar between groups. Subjects received a fixed show-up fee corresponding to 10 euros/hour of experiment. In addition, participants also gained a monetary bonus. This bonus was set at 10 euros for subjects in Group_-SR_ and Group_-S_ while it was variable from 0 to 20 euros according to the Group_-SRR_ performance (gain of 8 cents per successful trial in Block_-SRR_). Importantly, this bonus procedure in Block_-SRR_ was determined to match that obtained in the other groups; it corresponded to 10.4 ± 0.67 euros. A t-test revealed that the total ending remuneration was similar across the different Group_TYPES_ (t_(29)_ = 0.57; p = 0.57).

## Acknowledgements

We would like to thank Benvenuto Jacob and Julien Lambert for helping with the development of the task and Wanda Materne for assistance with data acquisition.

## Data availability

Data used in this study are available upon request (contact) and will be freely available at the time of publication via an open-access data sharing repository (https://osf.io).

